# Promoter boundaries for the *luxCDABE* and *betIBA-proXWV* operons in *Vibrio harveyi* defined by the method RAIL: Rapid Arbitrary PCR Insertion Libraries

**DOI:** 10.1101/227371

**Authors:** Christine M. Hustmyer, Chelsea A. Simpson, Stephen G. Olney, Matthew L. Bochman, Julia C. van Kessel

## Abstract

Experimental studies of transcriptional regulation in bacteria require the ability to precisely measure changes in gene expression, often accomplished through the use of reporter genes. However, the boundaries of promoter sequences required for transcription are often unknown, thus complicating construction of reporters and genetic analysis of transcriptional regulation. Here, we analyze reporter libraries to define the promoter boundaries of the *luxCDABE* bioluminescence operon and the *betIBA-proXWV* osmotic stress operon in *Vibrio harveyi*. We describe a new method called RAIL (<u>R</u>apid <u>A</u>rbitrary PCR <u>I</u>nsertion <u>L</u>ibraries) that combines the power of arbitrary PCR and isothermal DNA assembly to rapidly clone promoter fragments of various lengths upstream of reporter genes to generate large libraries. To demonstrate the versatility and efficiency of RAIL, we analyzed the promoters driving expression of the *luxCDABE* and *betIBA-proXWV* operons and created libraries of DNA fragments from these loci fused to fluorescent reporters. Using flow cytometry sorting and deep sequencing, we identified the DNA regions necessary and sufficient for maximum gene expression for each promoter. These analyses uncovered previously unknown regulatory sequences and validated known transcription factor binding sites. We applied this high-throughput method to *gfp, mCherry*, and *lacZ* reporters and multiple promoters in *V. harveyi*. We anticipate that the RAIL method will be easily applicable to other model systems for genetic, molecular, and cell biological applications.

**Importance:** Gene reporter constructs have long been essential tools for studying gene regulation in bacteria, particularly following the recent advent of fluorescent gene reporters. We developed a new method that enables efficient construction of promoter fusions to reporter genes to study gene regulation. We demonstrate the versatility of this technique in the model bacterium *Vibrio harveyi* by constructing promoter libraries for three bacterial promoters using three reporter genes. These libraries can be used to determine the DNA sequences required for gene expression, revealing regulatory elements in promoters. This method is applicable to various model systems and reporter genes for assaying gene expression.

## Introduction

Central to the study of bacterial physiology and development is the ability to monitor and quantify gene expression. Monitoring gene expression is greatly aided through the use of gene reporter fusions. Transcriptional and translational fusion constructs facilitate single-cell and population-wide gene expression investigations to study the influence of regulatory factors, perform genetic screens, and visualize protein localization patterns. Typically, such reporters are cloned downstream of regulatory promoters or genes of interest and introduced into a model bacterial system, either on replicating plasmids or integrated into the genome. Numerous reporter genes have traditionally been used to assay gene expression, such as *lux* (bacterial luciferase), *lacZ* (β-galactosidase), *phoA* (alkaline phosphatase), *bla* (β-lactamase), and *cat* (chloramphenicol acetyltransferase) (1, 2). However, the advent of more modern techniques has allowed for the use of fluorescent proteins such as green fluorescent protein (GFP) for these studies without the need for substrates or specialized media (1–4).

To adequately and efficiently study the expression pattern of a particular gene, the defined regulatory region controlling promoter activity must be known. The region upstream of the promoter driving *luxCDABE* transcription in *Vibrio harveyi* is an example of a locus with a large and undefined regulatory region, which has limited studies of gene regulation. This particular promoter is of interest because it drives expression of the bioluminescence genes with >100-fold increase in transcription and >1000-fold increase in bioluminescence production under activating conditions (*i.e*., quorum sensing) (5–10). It was previously suggested that the *lux* promoter requires ~400 bp upstream of the translation start site and ~60 bp downstream of the start codon for full activation of the *cat* reporter gene (9). The requirement for a large promoter region is due in part to the presence of seven binding sites for the transcription factor LuxR upstream of the primary transcription start site, each of which is necessary for maximal activation of the promoter (5, 6, 9). The ~400-bp region of P_*luxCDABE*_ is relatively large compared to some bacterial regulatory promoters (*e.g*., the *lac* promoter), but comparable in size to other promoters with evidence of cooperative binding between transcription factors and DNA looping (*e.g*., the *araBAD* promoter) (5, 11–13). Indeed, full activation of the *luxCDABE* promoter requires the transcription factor LuxR and nucleoid-associated protein Integration Host Factor (IHF), and DNA looping by IHF is proposed to drive interaction between LuxR and RNA polymerase for transcription activation (6).

Another *V. harveyi* operon that has an unknown promoter region is the *betIBA-proXWV* operon in *V. harveyi*. The *betIBA-proXWV* osmoregulation genes encode proteins required for the synthesis and transport of the osmoprotectant glycine betaine (14). These genes are auto-regulated by the BetI repressor and activated 3- to 10-fold by LuxR (14). Two LuxR DNA binding sites in the *betIBA-proXWV* promoter were identified by bioinformatics, and LuxR binding to these sites has been confirmed *in vitro* by electrophoretic mobility shift assays (14). In addition, ChIP-seq shows that LuxR binds with high affinity to two binding sites at this locus *in vivo* (5). For both the *luxCDABE* and *betIBA-proXWV* operons, the boundaries of the promoters are not defined, and thus, mechanistic studies of transcriptional regulation of these operons is limited.

Here, we describe a new method for rapidly generating reporter plasmids that we used to define promoter regions. The RAIL method (<u>R</u>apid <u>A</u>rbitrary PCR <u>I</u>nsertion <u>L</u>ibraries) exploits the power of arbitrary PCR and isothermal DNA assembly (IDA) to insert semi-randomized fragments of promoter DNA into reporter plasmids (15–17). Using RAIL, we generated libraries containing fragments of various lengths of the region upstream of the *luxCDABE* operon transcriptionally fused to *gfp*. We used flow cytometry sorting to screen the library of promoter fragments for reporter expression and next-generation sequencing to map the 3’ boundary of the *luxCDABE* promoter required for full activation. We also applied this method to two additional promoter regions in *V. harveyi* (*betIBA-proXWV* and *VIBHAR_06912*), and we demonstrated the versatility of the system by using two additional reporters, mCherry and β-galactosidase. This approach enabled us to identify the required regions for gene expression for multiple promoters and simultaneously produce usable gene reporter constructs. Our method should be widely applicable to any system for which gene reporters have been established and represents a simple and efficient technique to construct reporter fusions for molecular, genetic, and cell biology studies.

## Results

### Measuring transcription activation from the V. harveyi luxCDABE promoter using fluorescent reporter fusions

To study the mechanism of LuxR regulation of the *luxCDABE* promoter, we constructed four reporter plasmids containing various fragments of the *luxCDABE* locus transcriptionally fused to *gfp* using traditional cloning methods (Fig. 1A). Each plasmid contains the same 5’ end (~400 bp upstream of the *luxC* ORF), and the 3’ ends vary as follows: 1) 2 bp after the transcription start site at −26 (pJV369), 2) at the LuxC translation start site (pJV367), 3) 36 bp into the *luxC* ORF (pSO04), and 4) 407 bp into the *luxC* ORF (pJV365) (Fig. 1A). We first tested LuxR activation of these reporter plasmids in *Escherichia coli* because expression of *luxR* in *E. coli* is sufficient to drive high levels of transcription of the *luxCDABE* operon (5, 10, 18), and the use of *E. coli* is more efficient for transformation. Transcription activation of the *luxCDABE* promoter was assayed in *E. coli* strains containing a second plasmid either constitutively expressing *luxR* (pKM699) or an empty vector (pLAFR2). As shown previously, a plasmid containing the 3’ boundary 36 bp into the *luxC* ORF was highly expressed (Fig. 1B) (5). The strain containing pJV369 with the DNA fragment up to and including the transcription start site also displayed high levels of GFP. Activation was appreciably decreased (~7-fold) for the pJV367 strain containing the 5’ untranslated region (5’-UTR) but ending at the LuxC translation start site compared to the pSO04 strain (Fig. 1B). Also, the strain containing pJV365 with 407 bp of the *luxC* ORF was not activated above 2-fold (Fig. 1B). A similar trend was obtained when these constructs were conjugated into *V. harveyi* strains and the GFP expression was compared between wild-type and Δ*luxR* strains (Fig. S1B). From these data, we conclude that a promoter fragment ending 2 bp past the transcription start site is sufficient for activation.

**Figure 1.**
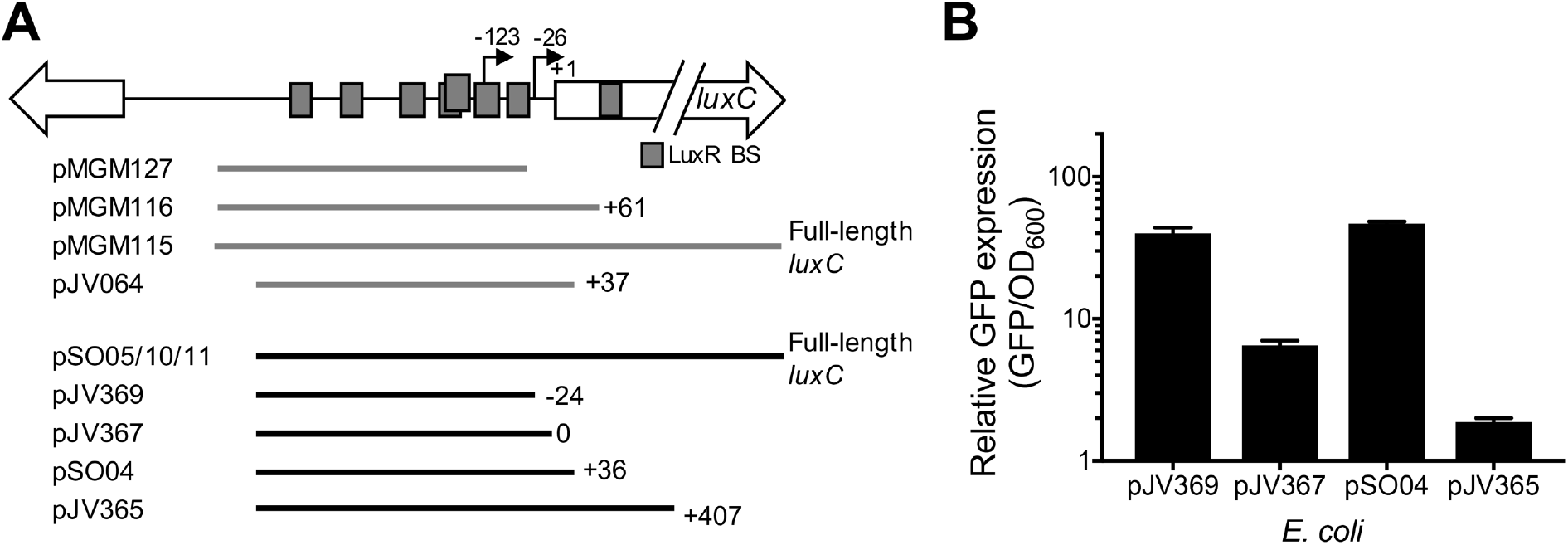
Promoter fusion plasmids for the *luxCDABE* genes. (A) Diagram of the regions of the *luxCDABE* promoter present in various plasmids listed, either from this study (black lines) or previous studies (gray lines) (5, 9). LuxR binding sites (LuxR BS) are shown as gray boxes. Transcription start sites are indicated by black arrows. The LuxC translation start site is shown as +1. Lengths of constructs are shown relative to the LuxC translation start site. (B) Relative GFP expression per OD_600_ (GFP/OD_600_) is shown for *E. coli* strains containing plasmids with varying *luxCDABE* promoter fragments fused to *gfp* as indicated in (A). The strains also contain either a plasmid constitutively expressing LuxR (pKM699) or an empty vector (pLAFR2). Relative expression was calculated by dividing the values for the pKM699-containing strain by the pLAFR2-containing strain.

### The RAIL method

Our observation that varying 3’ ends of the *luxCDABE* promoter greatly affected gene expression lead us to expand our analysis of the expression profile of promoter fusions across the entire locus. Therefore, we needed to construct numerous promoter fragments transcriptionally fused to a fluorescent reporter. Instead of constructing each of these plasmids individually, we designed a cloning technique combining the power of arbitrary PCR and IDA (a.k.a., Gibson assembly) (15–17). This method enabled us to simultaneously amplify fragments of varying lengths and clone them into a vector backbone to create a library in four simple steps (Fig. 2). First, arbitrary primers were used in a preliminary round of PCR in conjunction with a primer that specifically anneals to the promoter (Fig. 2, primer 1F). Four arbitrary primers were synthesized with eight sequential random nucleotides anchored at the 3’ end with two specific nucleotides: AA, TT, AT, or TA (Table S1). Each of these four primers also contains a linker at the 5’ end (Fig. 2, primer 1R). The first round of PCR produced a range of products that varied in length from 100 to >3000 bp and that appeared as faint smears of products as expected for random priming (Fig. 2). For some loci, no smear could be visualized by gel electrophoresis after the first round of PCR, but this did not impact the success of the second round of amplification.

**Figure 2.**
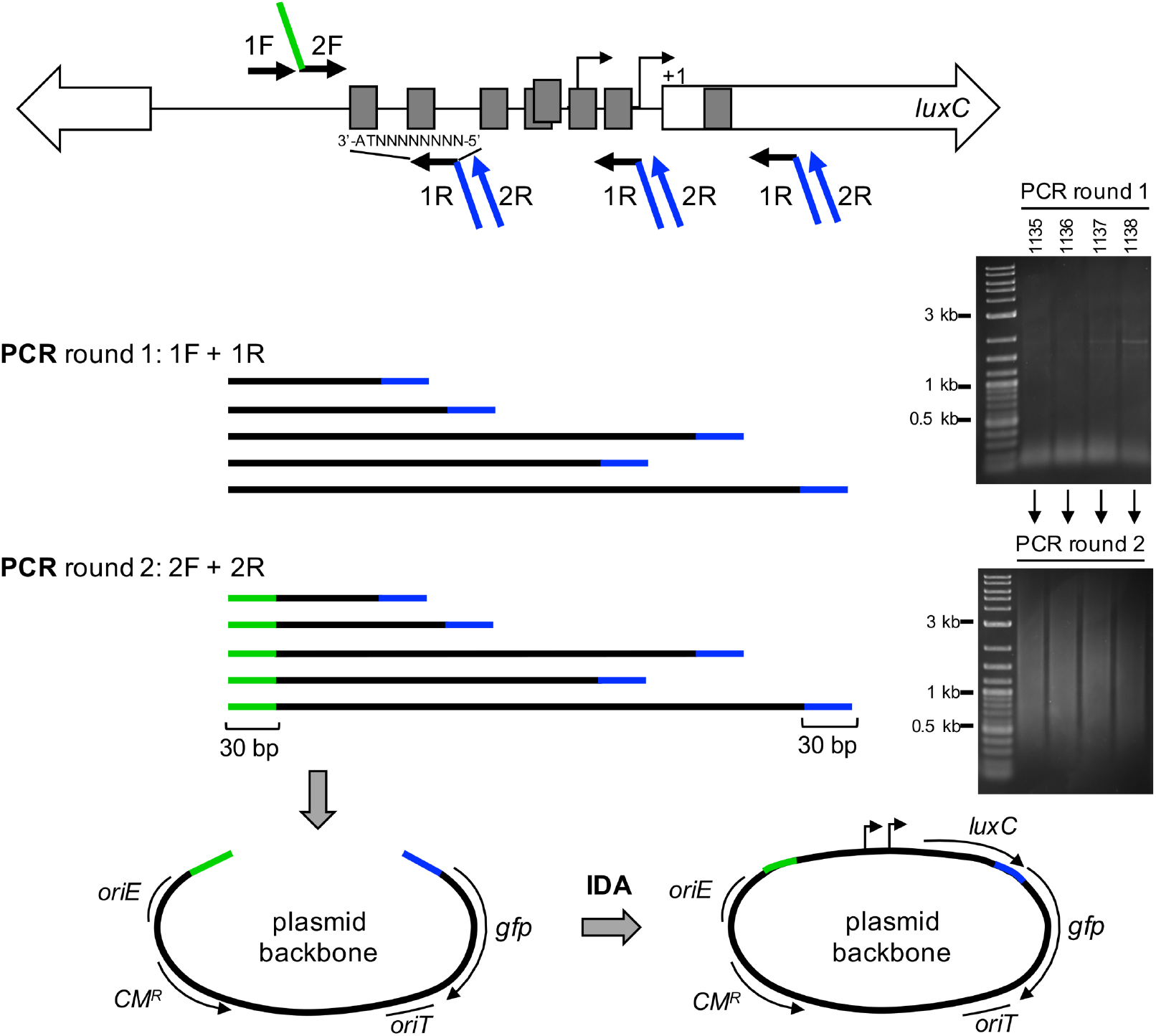
Schematic of the RAIL method for constructing libraries of promoter fusions. In PCR round 1, primers 1F and 1R were used to amplify a range of products specific to the *luxC* promoter. The 1R primers have eight random (‘N’) nucleotides incorporated, are anchored by two nucleotides (AA, AT, TA, or TT), and contain a linker (shown in blue). In PCR round 2, the 2F and 2R primers were used to further amplify and add a linker to the products. Primer 2F anneals just downstream of 1F and contains a linker (shown in green). Primer 2R anneals to the linker region of primer 1R. The linear plasmid backbone was prepared either by restriction digest or PCR. The library of products was inserted into the linear plasmid backbone via IDA using the homologous sequences present in the two linker regions (green and blue). The final plasmids contain fragments of the *luxC* promoter fused to the reporter. Gel images shown are examples of products from PCR rounds 1 and 2 for the *luxCDABE* locus using arbitrary R1 primers JCV1135-1138 and F1 primer SO71 in round 1 (Table S1). The products of the round 1 reactions were used in round 2 with primers SO72 and JCV1139 (Table S1).

In the second step, the products from round 1 were further amplified using a nested primer (primer 2F), and a linker was added with homology to the plasmid backbone (Fig. 2). The second round of PCR using these primers was performed with the products from round 1 as templates. Primer 2R anneals to the linker on primer 1R. This second step served to increase the amount of DNA product and to add a linker to the 5’ end. Each reaction in round 2 produced a smear of products that contained homology to the plasmid backbone at their 5’ and 3’ ends (Fig. 2). The smear of products can also be gel extracted to the desired size. In the third step, the plasmid backbone was PCR-amplified to form a linear product (Fig. 2). In the fourth and final step, IDA was performed to clone the promoter fragments into the plasmid backbone, and the mixture was transformed into *E. coli* to obtain isolated clones (Fig. 2).

### Defining the 3’ boundary of the luxCDABE operon using RAIL

We used the RAIL method to generate a large library of plasmids with promoter fragments fused to *gfp*. This library had fixed 5’-ends and varying 3’-ends generated by combining PCR products from four arbitrary primers, as shown in Figure 2, and inserts ranging from ~50 to >1,000 bp. We screened for *gfp* activation using fluorescence-activated cell sorting (FACS). The libraries were sorted by FACS into four groups: no GFP expression, low GFP expression, medium GFP expression, and high GFP expression (Fig. 3A). Illumina sequencing of the plasmid DNA from these pools enabled us to visualize the 3’ terminal end of the region cloned into the plasmid by graphing the location of the sequencing coverage (42 bp) and the 3’ terminal nucleotides (Fig. 3B, Fig. S2A). From these graphs, we pinpointed the boundary in the *luxCDABE* promoter required for maximum expression and showed the expression profile for promoter fragments across the entire locus (Fig. 3B, Fig. S2). The plasmids containing promoter fragments that terminated at nucleotide +129 (relative to +1, the start of the *luxC* ORF) were highly enriched in the ‘high expression’ pool, and plasmids in the ‘no expression’ pool were specifically de-enriched in this same location (Fig. 3B, Fig. S2A). A DNA fragment that terminates at +129 includes LuxR sites A, B, C, D, E, F, and G (6). The observation that LuxR site H was not included in this region of ‘high expression’ is consistent with previous findings that site H is non-essential for transcription activation at high cell density in *V. harveyi* (6). The ‘high expression’ pool had a clear 3’ boundary at +129, which is 16 bp upstream of LuxR site H. We conclude from these data that fragments with 3’ ends longer than +129 were decreased in reporter gene expression. There is also a clear edge where sequencing coverage drops off for the ‘high expression’ pool at −55 (Fig. 3B). However, the exact minimum boundary cannot be determined because we did not use every combination of anchor nucleotides in the arbitrary primers. The ‘medium expression’ pool contained sequences that terminated at +199, which is located 32 bp beyond LuxR site H (Fig. 3B). Plasmids with promoter fragments that extended throughout the *luxC* ORF past +199 had low levels of expression, whereas plasmids without GFP expression were limited to promoter regions upstream of −55 (Fig. 3B). We conclude that long promoter fragments decrease GFP expression and are not suitable reporter plasmids. We also conclude that plasmids containing promoter fragments shorter than −55 are not sufficient to activate transcription. Collectively, these data located the DNA region that is necessary and sufficient for maximal transcription activation (−393 to +129), validated previous findings that LuxR sites A through G are required for activation of *luxCDABE* (6), and defined the expression profile for the *luxCDABE* locus in the context of a transcriptional fusion to *gfp*.

**Figure 3.**
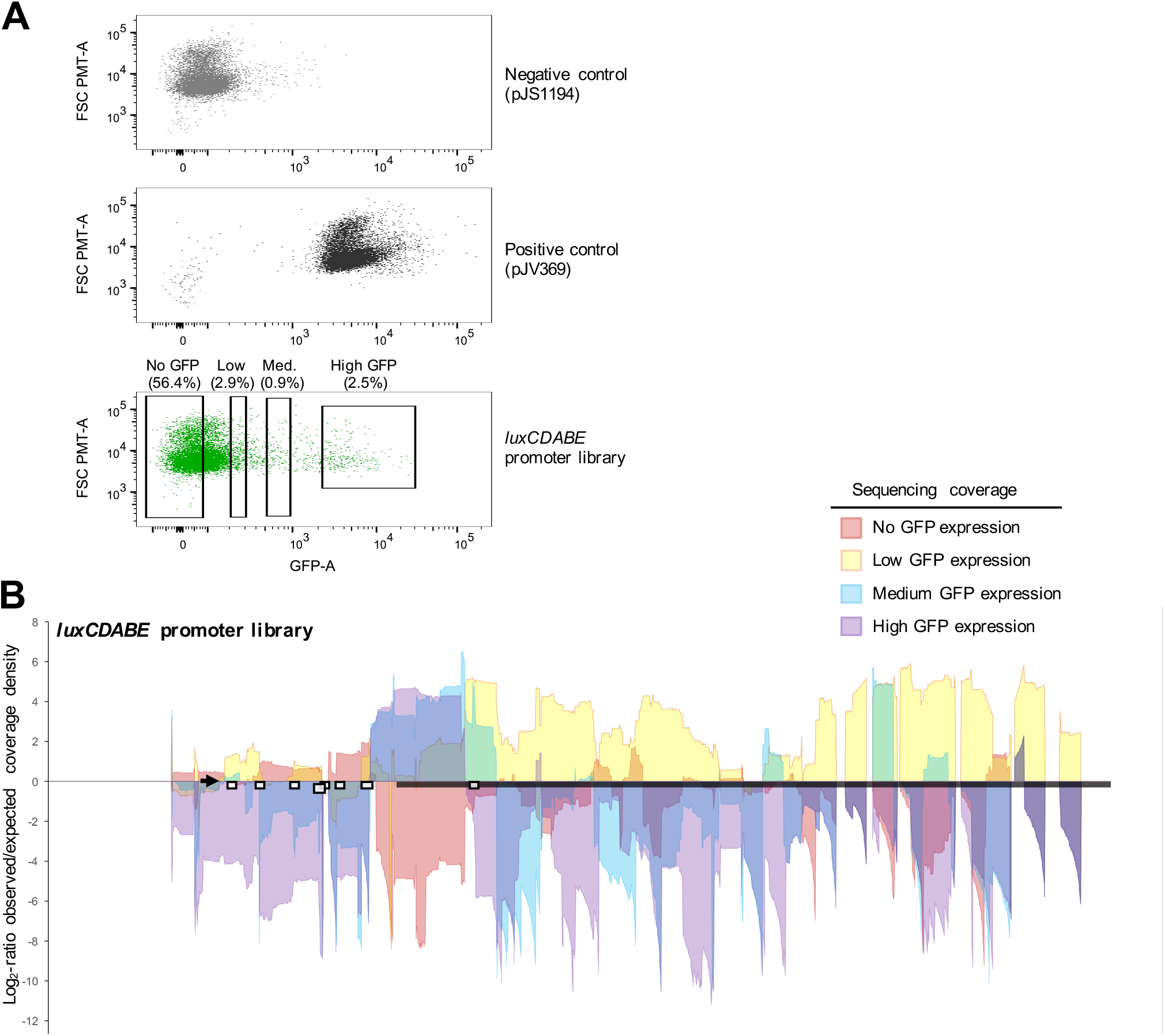
Flow cytometry sorting and next-generation sequencing of the P_*luxCDABE*_-*gfp* library. (A) FACS GFP expression data from three *E. coli* cultures: 1) a negative control strain containing an empty vector control (pJS1194) and a plasmid expressing LuxR (pKM699), 2) a positive control strain containing a P_*luxCDABE*_-*gfp* reporter plasmid (pJV369) and pKM699, and 3) the P_*luxCDABE*_ library of plasmids in *E. coli* containing pKM699. Gates (boxes) indicate the cells sorted (% of total population) into four bins: no GFP expression, low GFP expression, medium GFP expression, and high GFP expression. Data were presented using FlowJo software. (B) Genomic regions associated with differential expression of the *luxCDABE* locus. The nucleotide coverage of the reads (42 bp) is shown for different populations of cells with distinct levels of GFP reporter expression as indicated in the legend. Data are graphed as the nucleotide position (x-axis) versus the log_2_-ratio of observed coverage density divided by the expected coverage density (as determined by the read counts observed in the total library; y-axis). Areas with positive values in log2 observed/expected coverage densities indicate an enrichment of sequence reads in that region, and areas with negative values indicate lower than expected frequency reads. The thick black bar indicates the location of the *luxC* ORF. The locations of LuxR binding sites are indicated by white boxes. The locus to which the 2F primer anneals is indicated by an arrow.

### Defining the 3’ boundary of the betIBA-proXWV operon using RAIL

We next used the RAIL strategy to construct reporter clones for the *betIBA-proXWV* operon using a different fluorescent reporter, *mCherry*. We screened the promoter clones individually before using the high-throughput flow cytometry method to analyze the library. Approximately 40 plasmids were screened by restriction digest for inserts of varying sizes, and the inserts were sequenced to determine the size of the inserted region. We observed that plasmids containing regions shorter than the predicted transcription start sites did not show any activation compared to the empty vector control strain (Fig. 4B, pCH28 as an example). However, plasmids with larger regions that extended into the *betI* ORF were activated by LuxR, such as pCH50 and pCH72 (Fig. 4B). Plasmids containing the entire *betI* gene also did not display activation (Fig. 4B, pCH75).

**Figure 4.**
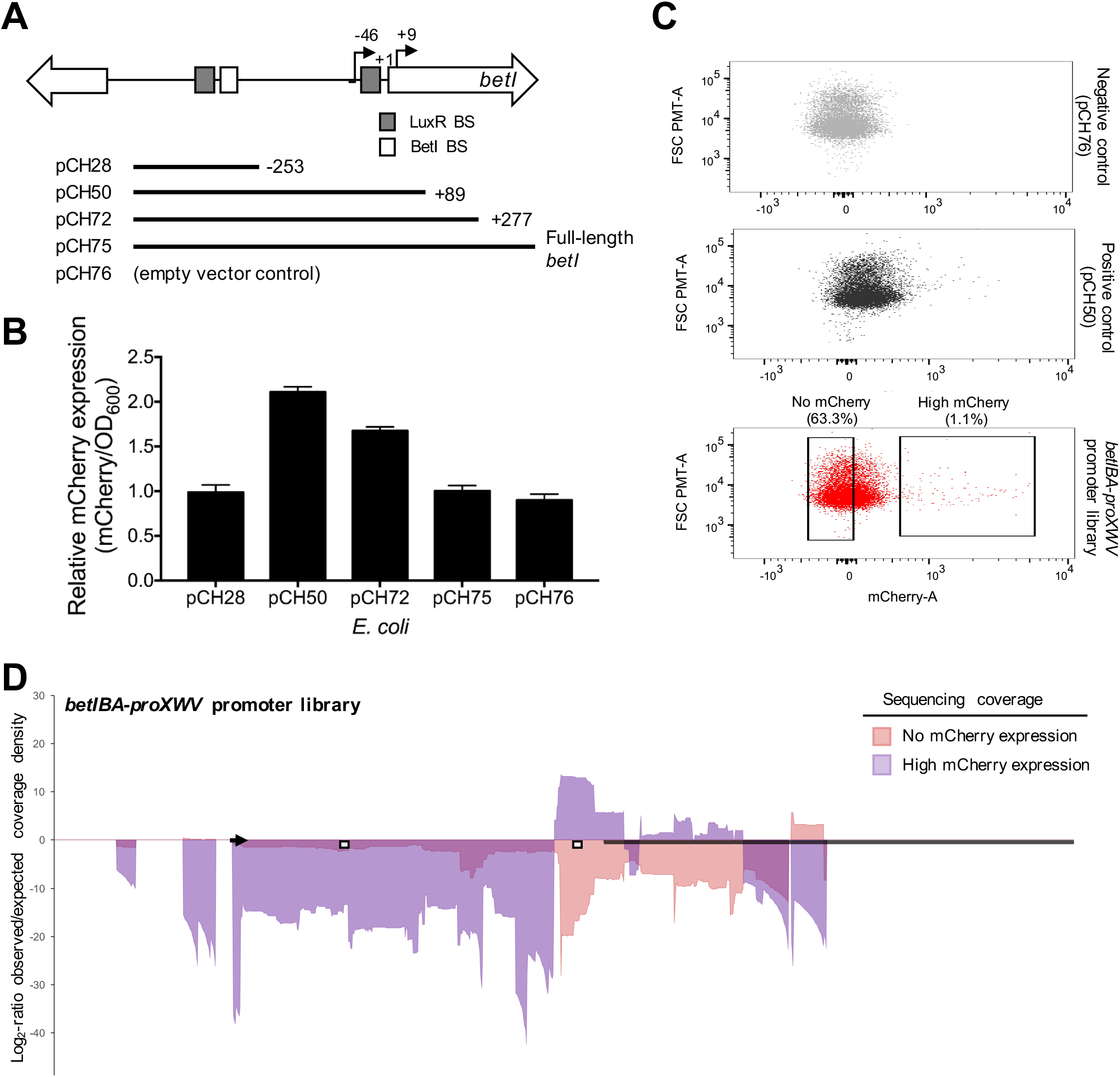
Expression data from the P_*betIBA-proXWV*_-*mCherry* library. (A) Diagram of the regions of the *betIBA-proXWV* promoter present in various plasmids. LuxR binding sites (LuxR BS) and the BetI binding site (BetI BS) are shown as gray and white boxes, respectively. Putative transcription start sites are indicated by black arrows. The BetI translation start site is shown as +1. Lengths of constructs are shown relative to the BetI translation start site. (B) Relative mCherry expression per OD_600_ (mCherry/OD_600_) is shown for *E. coli* strains as calculated by dividing the values for the pKM699 strain by the pLAFR2 strain. The strains contained plasmids with varying *betIBA-proXWV* promoter fragments fused to *mCherry* as indicated in (A). (C) FACS data showing mCherry expression for three *E. coli* cultures: 1) a negative control strain containing pCH76 and pKM699, 2) a positive control strain containing pCH50 and pKM699, and 3) the P_*betIBA-proXWV*_ library in *E. coli* containing pKM699. Gates indicate the cells sorted (% of total population) into two bins: no mCherry expression and high mCherry expression. (D) Genomic regions associated with differential expression of the *betIBA-proXWV* locus. The nucleotide coverage of the reads (42 bp) is shown for different populations of cells with distinct levels of or mCherry reporter expression as indicated in the legend. Data are graphed as the nucleotide position (x-axis) versus the log_2_-ratio of observed coverage density divided by the expected coverage density (as determined by the read counts observed in the total library; y-axis). Areas with positive values in log_2_ observed/expected coverage densities indicate an enrichment of sequence reads in that region, and areas with negative values indicate lower than expected frequency reads. The thick black bar indicates the location of the *luxC* ORF. The locations of LuxR and BetI binding sites are indicated by white boxes. The locus to which the 2F primer anneals is indicated by an arrow.

We synthesized a large library of *betIBA-proXWV* promoter fusions to *mCherry* using RAIL. It is important to note that only one arbitrary primer was used to generate this library, which limited the range of PCR products across the locus. This library of clones was sorted by FACS for those that maximally expressed *mCherry* (Fig. 4C), and the Illumina sequencing coverage and 3’ terminal nucleotides of the DNA in the two pools was graphed (Fig. 4D, Fig. S2B). Sequencing analyses revealed the minimum 3’ boundary for the *betIBA-proXWV* promoter to be at −13 (Fig. 4D, Fig. S2B; relative to +1, the start of the *betI* ORF), suggesting that the −46 transcription start site is the primary site for this locus. The ‘high expression’ pool contained plasmids with DNA fragments up through the first portion of the *betI* ORF at +25, which then tapered off (Fig. 4D, Fig. S2B). Plasmids with fragments that extended more than half-way through the *betI* gene displayed low or no expression. Collectively, these data showed that similarly to the *luxCDABE* locus, transcription reporters were functional if they contained DNA fragments past the 3’ boundary near the transcription start site. However, longer fragments extending into the ORF decreased reporter gene expression.

### Versatility of the RAIL method for cloning with other promoters and reporter genes

We also successfully used the RAIL technique to generate a promoter library using the *lacZ* reporter for another *V. harveyi* gene, *VIBHAR_06912*, which encodes a transcription factor. *VIBHAR_06912* expression is repressed by LuxR (19), and this is likely indirect repression because there are no detectable LuxR binding sites in this region (5). Using RAIL, multiple clones with varying promoter lengths were generated as transcriptional fusions to *lacZ*, and strains were assayed for β-galactosidase activity in *V. harveyi* (Fig. 5A). All of the plasmids with long promoter lengths were repressed by LuxR in the wild-type strain compared to the Δ*luxR* strain (Fig. 5B). Conversely, a plasmid with a short fragment (pJV342) showed the same level of β-galactosidase activity in the wild-type strain as in the Δ*luxR* strain (Fig. 5B). Thus, we conclude that we again generated functional promoter fusion plasmids for this promoter for future studies of gene expression and regulation of *VIBHAR_06912*.

**Figure 5.**
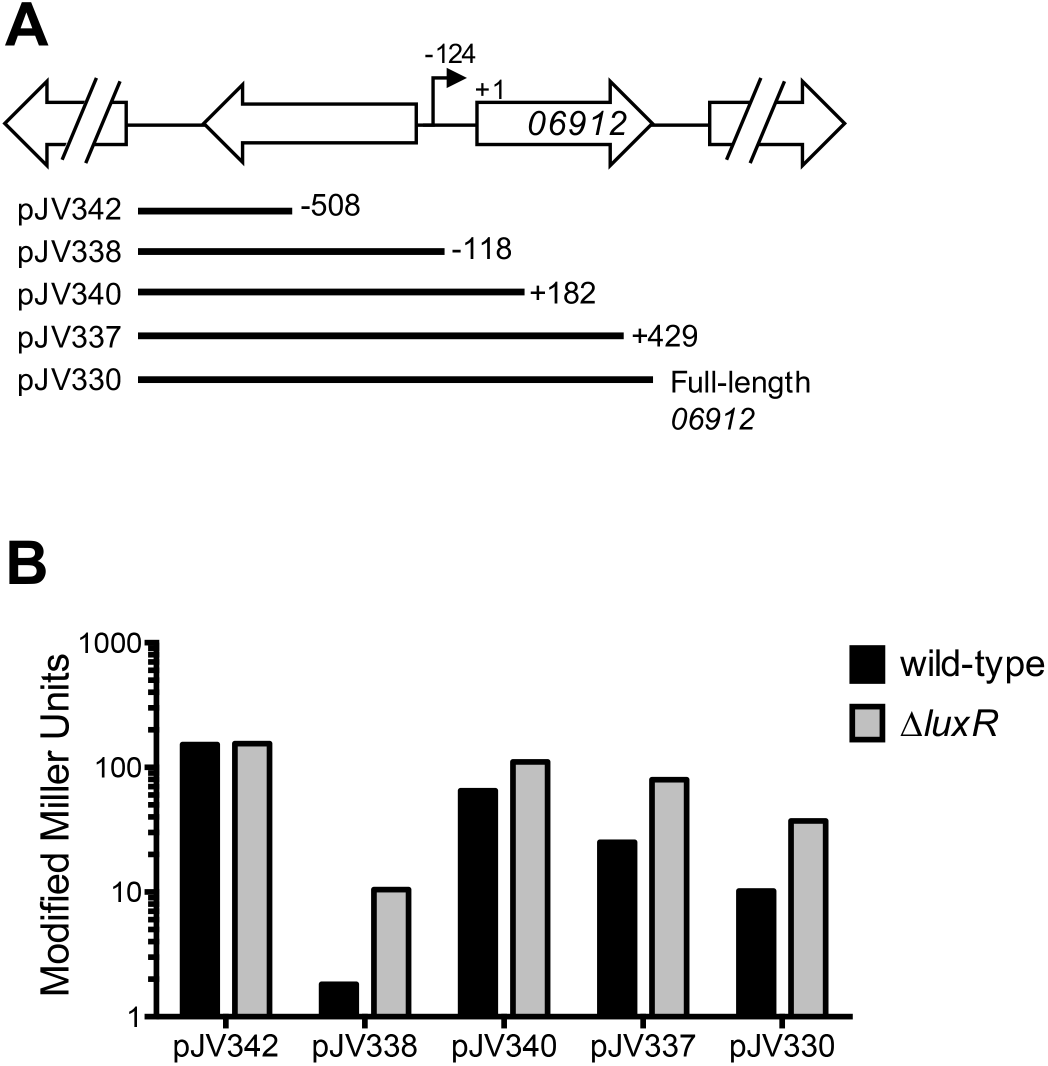
Promoter-reporter fusion plasmids for the *VIBHAR_06912* promoter. (A) Diagram of the regions of the *VIBHAR_06912* promoter present in various plasmids. The putative transcription start site is indicated by a black arrow. The VIBHAR_06912 translation start site is shown as +1. Lengths of constructs are shown relative to the VIBHAR_06912 translation start site. (B) Modified Miller units are shown for wild-type (BB120) and Δ*luxR* (KM669) V. harveyi strains containing various plasmids as indicated in (A). Data shown are representative of four independent biological experiments.

### Reporter gene affects measurement of gene expression

We noted that for each of the three promoters we studied, plasmid constructs that contained promoter regions that extended into the ORF of the first gene had variable levels of expression. For example, the pJV365 plasmid that included 407 bp of the *luxC* gene only expressed GFP ~2-fold more in the presence of LuxR than in its absence (Fig. 1B). This is in contrast to plasmid pMGM115 from the Miyamoto *et al*. study that contains the full *luxC* ORF and displays maximal activation of the *cat* gene (~50-fold more than truncated promoters) (9). To examine these contradictory results further, we constructed plasmids containing the entire *luxC* gene and its promoter region driving expression of *gfp, lacZ*, or *mCherry* (Fig. 1A, 6A). These constructs contained the intragenic region between *luxC* and *luxD* (15 bp), and the reporter gene was cloned in place of the *luxD* ORF (Fig. 6A). We observed that the *lacZ* and *mCherry* plasmids were activated 16- to 20-fold, whereas the *gfp* construct was only activated 1.6-fold by LuxR in *E. coli* (Fig. 6B). The *gfp* (pSO05) and *mCherry* (pSO11) plasmids had similar levels of activation when the plasmids were introduced into wild-type *V. harveyi*, though neither were expressed maximally (Fig. S1A).

**Figure 6.**
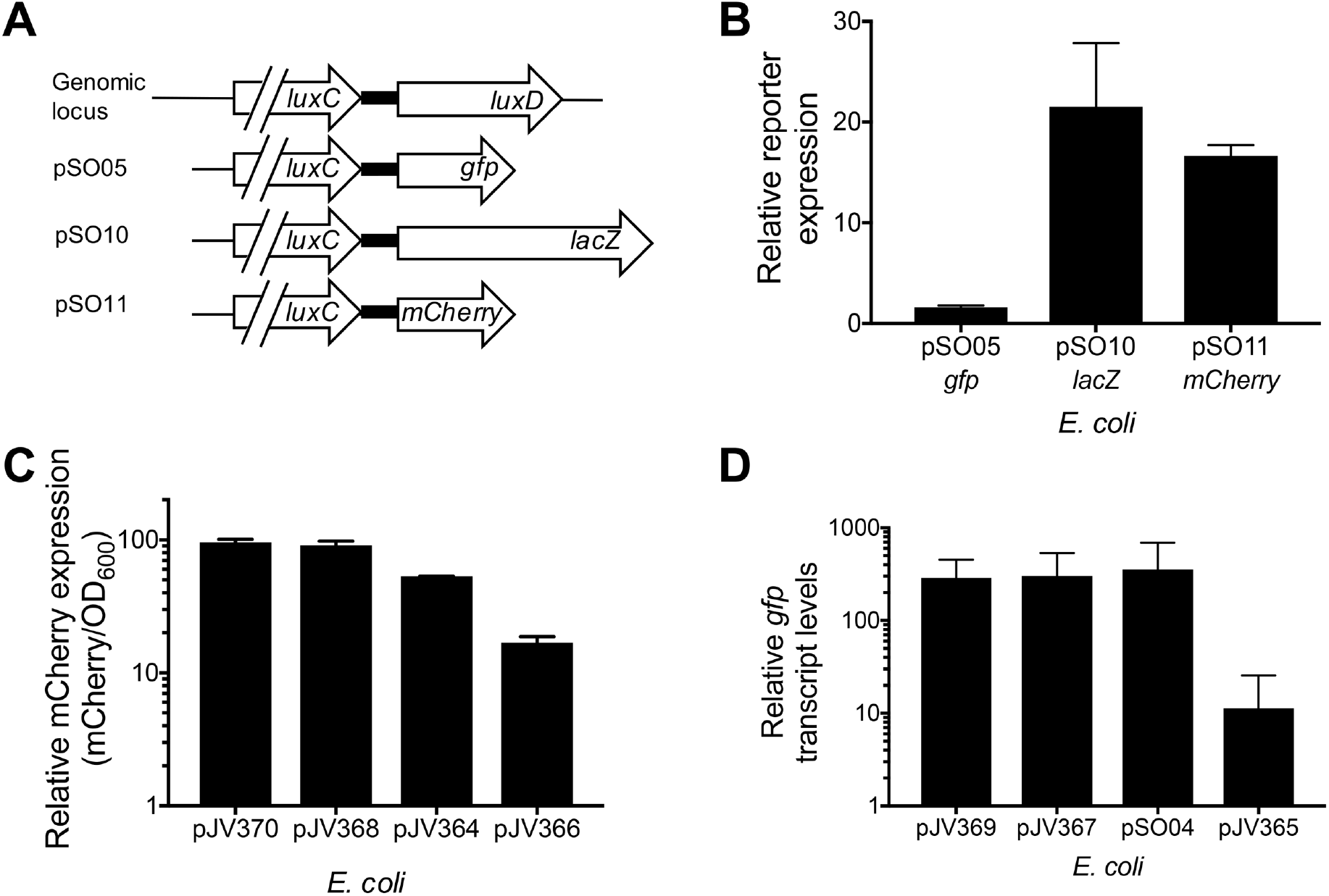
Measuring transcription of plasmids with long promoter regions fused to reporters. (A) Diagram of plasmids containing *luxCDABE* promoters and the *luxC* ORF fused to *gfp* (pSO05), *lacZ* (pSO10), or *mCherry* (pSO11). Each construct contains the 15-bp sequence between *luxC* and *luxD* as shown. (B) Relative expression of reporters (GFP, mCherry, and LacZ) is shown for *E. coli* strains containing either a plasmid constitutively expressing LuxR (pKM699) or an empty vector (pLAFR2). Relative expression was calculated by dividing the values for the pKM699-containing strain by the pLAFR2-containing strain. LacZ expression was determined by modified Miller assays, and GFP and mCherry expression were assayed using a plate reader. (C) Relative expression of mCherry (mCherry/OD_600_) is shown for *E. coli* strains containing either pKM699 or pLAFR2, calculated as described in (B). (D) Relative transcript levels of *gfp* determined by qRT-PCR for *E. coli* strains containing either pKM699 or pLAFR2, calculated as described in (B).

We hypothesized that the observed decrease in activation with longer fragments might be due to instability of the transcript when the *luxC* ORF is present upstream of the *gfp* reporter. Thus, we constructed *mCherry* reporter plasmids containing the same four *luxCDABE* promoter fragments that were fused to *gfp* in Figure 1A and assayed these in *E. coli* (Fig. 6C, Fig. S1A). We verified that the shortest region tested (2 bp past the primary transcription start site) was sufficient for activation, and there were minimal differences in expression with constructs containing the three shortest promoter fragments with all three being activated >50-fold (Fig. S1C). However, as seen with GFP, plasmids with longer promoter fragments (*e.g*., pJV366 with 407 bp of the *luxC* ORF) yielded a lower level of mCherry expression (Fig. 6C, activated 17-fold), which was similar to the construct containing the entire *luxC* ORF (Fig. 6B, pSO11, activated 17-fold). The same trend was observed in *V. harveyi* for these mCherry plasmids (Fig. S1B). Thus, we conclude that constructs containing long fragments of the *luxC* ORF indeed decrease expression of downstream reporters, for some of which large decreases occur (*i.e., gfp*). This result is not observed with expression of the *luxCDABE* operon *in vivo*; the expression levels of each of the five genes in the operon are similar and do not differ by more than 2-fold from one another (19).

To examine these results, we measured transcript levels of *gfp* for several *P_luxC_* reporter plasmids in *E. coli*. The relative transcript levels of *gfp* from qRT-PCR measurements were high for the three plasmids containing short regions of the *luxCDABE* promoter, but as seen with GFP expression measurements, levels of *gfp* transcripts dropped ~25-fold from the pJV365 plasmid containing 407 bp into the *luxC* ORF compared to pJV369 (Fig. 6D). Thus, we conclude that the decrease in GFP expression is due to a decrease in transcript levels, which may be caused either by transcript instability or a decrease in transcription initiation or elongation in plasmids with long promoter fragments (*e.g*., pJV365). We did not observe a decrease in *gfp* transcript levels with pJV367 as observed with GFP expression (Fig. 1B), suggesting that the decrease in GFP expression may be due to constraints at the post-transcriptional or translational level. These results indicate that testing multiple promoter fusions is beneficial for identifying a promoter-reporter fusion that functions *in vivo* to mimic expression from the native locus.

## Discussion

We have developed the RAIL method for rapid construction of promoter fusion plasmids and demonstrated that this approach can be applied to multiple promoters and reporter genes. The RAIL strategy can be used to quickly generate a few reporters or to create large libraries of promoter fusions for high-throughput analysis of the regions that drive transcription activation. The method requires simple cloning steps, and once the system is designed for a particular plasmid backbone, only two locus-specific primers are needed. For our plasmid backbone, we designed arbitrary primer sets for creating fusions to *gfp, mCherry*, and *lacZ* that can be used with any gene locus (Table S1), and these primers can be easily modified for use in any plasmid with a reporter gene.

Our library sets revealed several important findings with regard to the expression profiles for the *luxCDABE* and *betIBA-proXWV* promoters. First, we validated previous work describing the requirement for LuxR binding sites in these promoters (5, 6, 9, 14). Second, we identified the promoter region that is required for high levels of transcription activation for these two promoters. We did not resolve the 3’ boundary to a specific nucleotide locus in these experiments because we did not use every combination of anchor nucleotides in the arbitrary primers and restricted our analysis to combinations of A and T pairs. However, with this resolution we clearly found a marked difference in plasmids containing various fragments of the promoters such that we could identify the largest region necessary and sufficient for maximum gene expression. Smaller fragments may be sufficient to drive the same level of gene expression, which can be tested with the full series of anchor nucleotides in the arbitrary primers. Further, future studies could use the same approach to map the 5’ boundary of these two promoters, which is a separate but intriguing question.

Third, our data conclusively demonstrate that there is no requirement for the region downstream of the transcription start site for full activation of the *luxCDABE* promoter. This finding is important because a previous study by Miyamoto *et al*. also tested promoter regions for *luxCDABE* via a *cat* promoter (9). Among the various constructs tested in that study, the pMGM127 plasmid contains a region truncated slightly upstream of the −26 transcription start site (the specific 3’ end is undefined in the article) and the pMGM116 plasmid includes a 3’ end at +61 relative to the *luxC* start codon (Fig. 1A). The shorter promoter in pMGM127 shows no transcription activation, whereas the longer promoter in pMGM116 had full activation of the *cat* reporter (9). These data and other observations have led to an anecdotal hypothesis in the field that there is an element downstream of the transcription start site that is required for full activation of the *luxCDABE* promoter. Our data refute this hypothesis because the pJV369 plasmid does not include the 5’-UTR and is maximally activated in both *E. coli* and *V. harveyi*.

Finally, our analysis of various promoter-reporter fusion plasmids demonstrated that not all reporter fusions are created equal and suggests that testing various reporter constructs for each gene of interest is beneficial to finding the optimal reporter for downstream assays. We noted that plasmid constructs with long fragments of the *luxCDABE* and *betIBA-proXWV* promoters that included sections of the first ORF in the operon were substantially decreased in expression, and we showed that this is effective at the transcript level for *luxC-gfp* fusions (Fig. 6D). However, we also noted that the strains containing the pJV367 plasmid that had a decrease in GFP fluorescence did not exhibit a decrease in *gfp* transcript levels (Fig. 6C). This result implies that the 7-fold decrease in GFP fluorescence is due to post-transcriptional or translational effects, such as mRNA secondary structure that may block translation initiation.

There are at least two possible reasons why plasmids containing long fragments of the *luxCDABE* promoter have decreased transcript levels. One possibility is that transcripts generated with fragments of the *luxCDABE* operon fused to the *gfp* gene may fold into unstable secondary structure and be subject to degradation. However, we suspect that this explanation is unlikely to be the cause of low expression for every plasmid with a fragment longer than +129, as we would predict that at least some would be stable. A second possibility is that LuxR binding to site H is acting as a roadblock to transcription elongation, which results in the abrupt drop in GFP expression after site H. Previously, we showed that scrambling site H does not decrease LuxR activation of β-galactosidase expression in a *luxC-lacZ* reporter plasmid under conditions in which LuxR is maximally expressed at high cell densities in *V. harveyi* (6). However, the results of our expression profiling experiment in *E. coli* with the *luxCDABE* promoter library suggests that plasmids that contain LuxR site H have decreased levels of transcription activation and are strictly in the ‘low GFP expression’ pool (Fig. 3B). LuxR has an extremely high affinity for site H with a *K_d_* of 0.6 nM, one of the tightest LuxR binding affinities in the genome (5). Thus, it is curious why LuxR binds at this locus with no apparent activation defect when tested at high cell density in *V. harveyi*.

Protein roadblocks have been described in bacteria and eukaryotes that hinder transcription, and elongation factors aid in transcription elongation through these roadblocks by various mechanisms (*e.g*., Mfd in *E. coli*) (20). In addition, when multiple RNAP molecules are initiated from the same promoter, these trailing RNAP complexes can “push” a stalled RNAP through a roadblock (21). Thus, it is possible that higher levels of transcription initiation of the *luxCDABE* promoter in *V. harveyi* at high cell densities drive transcription elongation through site H, whereas lower levels of LuxR at low cell densities in *V. harveyi* or in our synthetic *E. coli* system are not sufficient to push through the LuxR site H roadblock. LuxR concentrations are low in the cell at low cell densities, and thus, the relatively few LuxR molecules likely bind to the highest affinity sites, such as site H in the *luxC* ORF. As cells grow to high cell densities, LuxR levels accumulate (19, 22, 23) and enable LuxR binding to other sites, which drives high levels of transcription initiation and may relieve binding of LuxR to site H to allow RNA polymerase elongation. Alternatively, the roadblock might be relieved by restructuring of the DNA architecture at the locus. Because we have already shown that IHF binds to multiple places at the *luxCDABE* region and its binding is positively cooperative with LuxR, this DNA bending may play a role in removing transcription roadblocks. We also observed a sharp difference between the ‘medium expression’ pool and ‘low expression’ pool just downstream of the LuxR site H (Fig. 3B), suggesting that there may be yet another roadblock in this region. Future studies should elucidate the role of LuxR binding sites within ORFs in *V. harveyi*, which are observed throughout the genome (5).

In conclusion, the RAIL method offers a rapid and efficient method to obtain libraries of reporter fusions that can be used for various studies of gene expression and regulation. Often in bacterial genetics, researchers attempt to create promoter fusions by cloning a reporter gene in place of the translation start site, and this would have yielded suboptimal reporters for the *luxCDABE* promoter. Anecdotally, and as we experienced with the *betIBA-proXWV* and *luxCDABE* promoters, one often needs to construct multiple reporter fusions to identify a promoter region that drives gene expression mimicking native locus gene expression. Thus, our method is more efficient by generating numerous clones in a single cloning experiment. We envision use of the RAIL method for numerous other purposes, such as creating functional GFP protein fusions for studying protein localization, identifying cis-regulatory sequences in promoters (*e.g*., protein binding sequences), inserting affinity tags for purification strategies, and identifying highly expressed soluble constructs for protein purification. Finally, this method should be applicable to any model organism for which genetic cloning techniques have been established.

## Materials and Methods

### Bacterial strains and media

*E. coli* strains S17-1λpir, DH10B, and derivatives (Table S2) were used for cloning and *in vivo* assays. *E. coli* strains were grown shaking at 275 RPM at 37°C in lysogeny broth (LB), augmented with 10 μg/mL chloramphenicol and 10 μg/mL tetracycline when required. The *V. harveyi* BB120 is strain ATCC BAA-1116, which was recently reassigned to *Vibrio campbellii* (24). It is referred to as *V. harveyi* throughout this manuscript for consistency with previous literature. BB120 and derivatives (Table S2) were grown at 30°C shaking at 275 RPM in LB Marine (LM) medium supplemented with 10 μg/mL chloramphenicol when required. LM is prepared similarly to LB (10 g tryptone, 5 g yeast extract) but with 20 g NaCl instead of 10 g used in LB.

### Molecular methods

Oligonucleotides (Table S1) were purchased from Integrated DNA Technologies. All PCR reactions were performed using Phusion HF polymerase (New England BioLabs) or iProof polymerase (BioRad). Restriction enzymes, enzymes for isothermal DNA assembly (15), and dNTPs were obtained from New England BioLabs. DNA samples were visualized on 1% agarose gels. Standard cloning methods and primers for the single plasmid constructs listed in Table S3 are available upon request. Standard sequencing of single plasmid constructs was conducted by ACGT, Inc. and Eurofins Genomics. To measure the expression levels of fluorophore reporter plasmids, *E. coli* and *V. harveyi* strains were grown overnight at 30°C shaking at 275 RPM. Strains were diluted 100-fold in growth media and selective antibiotics in 96-well plates (black with clear bottom), covered with microporous sealing tape (USA Scientific), and incubated shaking at 30°C at 275 RPM for 16-18 h. Fluorescence and OD_600_ from strains expressing *mCherry* and *gfp* were measured using either a BioTek Synergy H1 or Cytation plate reader. Miller assays were conducted as previously described (6). RNA extraction and qRT-PCR were performed and analyzed as described (14) with primers listed in Table S1 on a StepOne Plus Real-Time PCR machine (Applied Biosystems). Transcript levels were normalized to the level of expression of the internal standard *recA*, and the standard curve method was used for data analysis. The error bars on graphs represent the standard deviations of measurements for triplicate biological samples.

### RAIL: construction of promoter libraries by arbitrary PCR

The arbitrary PCR method was adapted from Schmidt *et al*. (25) with several modifications. The first round of PCR was conducted using two primers: a forward primer specific to the promoter of interest and a reverse primer for random DNA amplification (Fig. 2, primers 1F and 1R, respectively). The 1R primer includes a priming sequence, followed by eight random nucleotides (‘N’), and terminating in two defined nucleotide anchors, either AT, TA, TT, or AA (Table S1). For PCR round 1, ~10-100 ng/μl of genomic DNA from *V. harveyi* BB120 or a plasmid containing the region of interest was added as the template. The reaction included 200 μM dNTPs, 250 μM primers, 5% DMSO, 0.5 μl Phusion polymerase, and 1X Phusion buffer. Cycling parameters were as follows and as previously published (25): an initial denaturation at 95°C for 5 min, then 5 cycles of 95°C for 30 s, 25°C for 30 s, and 72°C for 2.5 min, followed by 30 cycles of 95°C for 30 s, 50°C for 30 s, and 72°C for 2.5 min, and a final extension step of 72°C for 10 min. PCRs were purified using the GeneJet PCR Purification Kit (Thermo Scientific) and eluted in 30-50 μl of elution buffer. The second round of PCR used primers 2F and 2R (Fig. 2). The forward primer (2F) included 30 nt homology to the plasmid backbone for IDA and a sequence specific to the promoter of interest that is nested downstream of the 1F primer. The reverse primer (2R) included the priming sequence that is identical to that of primer 1R. To perform PCR round 2, 5 μl of the purified DNA from round 1 was used as the template. These reactions also included 200 μM dNTPs, 250 μM primers, 5% DMSO, 0.5 μl Phusion polymerase and 1X Phusion buffer. Cycling parameters were as follows: an initial denaturation at 95°C for 5 min, then 35 cycles of 95°C for 30 s, 50°C for 30 s, and 72°C for 2.5 min, and a final extension step of 72°C for 10 min. PCR products were separated by agarose gel electrophoresis, visualized by UV transillumination, and the products were gel extracted to desired target size using a GeneJet Gel Extraction Kit (Thermo Scientific).

Cloning of arbitrary PCR inserts into the plasmid backbone was performed using IDA as described (15). Library inserts were incubated in IDA reactions with 100 ng of plasmid backbone, and these reactions were transformed into electrocompetent *E. coli* Electromax DH10B cells (ThermoFisher) and plated on media with selective antibiotics. DNA from individual colonies was first screened by restriction digest and sequenced to confirm that inserts of the desired size were incorporated. For generation of libraries for sorting, >50,000 colonies were collected from plates, mixed in LB selective media, and the culture stored at −80°C. DNA extracted from this library was transformed into electrocompetent *E. coli* S17-1λpir cells containing a plasmid expressing *luxR* (pKM699). After this second transformation step, >50,000 colonies were collected from plates, mixed in LB selective media, and the culture stored at - 80°C.

### Flow cytometry sorting

A BD FACSAria II was used to sort *E. coli* cells based on expression of the fluorescent reporter. In preparation for the sort, the library culture from the frozen stock was grown in 50 mL of selective medium at 30°C shaking to OD_600_ = 0.5. DNA was extracted from this culture as the ‘input’ DNA sample for sequencing. Control cultures were used to set the sorting gates. The positive controls for maximal expression were strains containing pKM699 (expressing *luxR*) and a plasmid construct that demonstrated high levels of expression for either P_*luxC*_ (pJV369, P_*luxC*_-*gfp* positive control plasmid) or P_*betI*_ (pCH50, P_*betI*_-*mCherry* positive control plasmid). The negative controls were strains containing pKM699 and pJS1194 (*gfp* negative control plasmid) or pCH76 (*mCherry* negative control plasmid), which lack promoters in front of *gfp* or *mCherry*, respectively. The library culture was sorted into bins with >10,000 cells per bin. For the P_*luxC*_ sort, there were four bins: no GFP expression, low GFP expression, medium GFP expression, or high GFP expression. For the P_*betI*_ sort, there were two bins: high mCherry expression or no mCherry expression. The sorted cell cultures were incubated in 5 mL of selective medium shaking at 30°C at 275 RPM and grown to stationary phase. Cultures were stored at −80°C, and the DNA was extracted for sequencing using the GeneJet Miniprep kit (Thermo Scientific).

### Illumina library preparation and sequencing

DNA extracted from all sorted samples and input controls was purified over a Performa DTR gel filtration cartridge (Edge Biosystems). Next, the DNA was sheared using a Covaris S220 in 6 × 16 mm microtubes to average sizes of 400 bp and analyzed on an Agilent 2200 TapeStation using D1000 ScreenTape. The sheared DNA samples (1 μg) were each treated with terminal deoxynucleotidyl transferase (TdT) in a tailing reaction using TdT (Promega) and a mixture of dCTP and ddCTP (475 μM and 25 μM final concentrations, respectively) to generate a poly-C tail. The reactions were incubated at 37°C for 1 h, heat-inactivated at 75°C for 20 min, and cleaned over a Performa DTR gel filtration cartridge. Next, two rounds of PCR were performed to attach sequences for Illumina sequencing (Table S1). In round 1, a C-tail specific primer olj376 and a gene-specific primer (CH060 for the *gfp* library, and CH063 for the *mCherry* library) were used to amplify the promoter fragments that were cloned into the plasmid during the RAIL method. This PCR reaction used Taq polymerase (NEB), with the following cycling conditions: an initial denaturation at 95°C for 2 min, then 24 cycles of 95°C for 30 s, 58°C for 30 s, and 72°C for 2 min, and a final extension step of 72°C for 2 min. The round 2 PCR used a nested gene specific primer (CH061 for the *gfp* library, and CH064 for the *mCherry* library) to provide added specificity and also to append the linker sequence needed for Illumina sequencing. The second primer in the reaction contained different barcodes for each sample to enable the libraries to be pooled and sequenced simultaneously (Table S1; BC37-44). The round 2 PCRs were performed using Taq polymerase, and cycling as follows: an initial denaturation at 95°C for 2 min, then 12 cycles of 95°C for 30 s, 52°C for 30 s, and 72°C for 2 min, and a final extension step of 72°C for 2 min. These final PCR products were examined by DNA gel electrophoresis, at which point a smear of products was visible on the gel.

Sequencing was performed on a NextSeq 500 using a NextSeq 75 reagent kit using 42bp × 42bp paired-end run parameters and gene specific primers (CH062 for the *gfp* library, and CH065 for the *mCherry* library). Reads were checked with FastQC (v0.11.5; Available online at: http://www.bioinformatics.babraham.ac.uk/projects/fastqc) to ensure that the quality of the data was high and that there were no noteworthy artifacts associated with the reads. The R1 read from paired-end Illumina NextSeq reads were quality and adapter trimmed with Trimmomatic (v0.33) (26) using the following parameters: ILLUMINACLIP:<adapters>:3:20:6 LEADING:3 TRAILING:3 SLIDINGWINDOW:4:20 MINLEN:25. Reads were high quality with 80-85% of the reads surviving trimming. Reads were mapped against the *Vibrio campbellii* ATCC BAA-1116 genome (including molecules NC_009777, NC_009783, and NC_009784) using bowtie 2.3.2 (27) and visualized using JBrowse (version 1.10.12) to analyze the alignment to the *betI* and *luxC* gene regions. Analysis of the promoter-seq data hinged on contrasting high, medium, low, and no expressing cells to the null distribution. The null distribution was determined by sequencing cells collected without any sorting applied. The reads associated with the null, high, medium, low, and no expression datasets for the *luxC* library (or null, high, and no expression datasets for the *betI* library) were normalized by transforming the data into the fraction of bases covered, which was defined as the depth of coverage at a particular base divided by the total depth of coverage over the particular promoter region (*betI:* bases 1361141-1362440 on NC_009784; *luxC*: bases 1424774-1426907 on NC_009784). Analysis was performed by plotting the log_2_ ratio of the observed fraction of bases covered (for high, medium, low or no expression cell collections) over the expected (null) distribution.

## Acknowledgments

We thank Ankur Dalia for assistance with the design of the Illumina sequencing protocol and helpful discussions. We also thank James Ford and Douglas Rusch from the Indiana University Center for Genomics and Bioinformatics for Illumina sequencing experiments, bioinformatics analysis, and assistance with graphing the results. We also wish to acknowledge Christiane Hassel from the Indiana University Flow Cytometry Core Facility for assistance with FACSAria II analysis and sorting. We appreciate the use of Jason Tennessen’s BioTek Cytation plate reader, and we thank Ryan Chaparian and Blake Petersen for assistance with experiments. This work was supported by the National Institutes of Health (NIH) Grant 1R35GM124698-01 to JVK and start-up funds from Indiana University to JVK and MLB.

